# Calcium buffers and L-type calcium channels as modulators of cardiac subcellular alternans

**DOI:** 10.1101/683011

**Authors:** Yi Ming Lai, Stephen Coombes, Rüdiger Thul

## Abstract

In cardiac myocytes, calcium cycling links the dynamics of the membrane potential to the activation of the contractile filaments. Perturbations of the calcium signalling toolkit have been demonstrated to disrupt this connection and lead to numerous pathologies including cardiac alternans. This rhythm disturbance is characterised by alternations in the membrane potential and the intracellular calcium concentration, which in turn can lead to sudden cardiac death. In the present computational study, we make further inroads into understanding this severe condition by investigating the impact of calcium buffers and L-type calcium channels on the formation of subcellular calcium alternans when calcium diffusion in the sarcoplasmic reticulum is strong. Through numerical simulations of a two dimensional network of calcium release units, we show that increasing calcium entry is proarrhythmogenic and that this is modulated by the calcium-dependent inactivation of the L-type calcium channel. We also find that while calcium buffers can exert a stabilising force and abolish subcellular Ca^2+^ alternans, they can significantly shape the spatial patterning of subcellular calcium alternans. Taken together, our results demonstrate that subcellular calcium alternans can emerge via various routes and that calcium diffusion in the sarcoplasmic reticulum critically determines their spatial patterns.

## 1. Intoduction

Cardiac arrhythmias constitute a leading public health problem and cause most cases of sudden cardiac death. In the US alone, sudden cardiac death accounts for approximately 300, 000 – 450, 000 lives every year [1]. Among the many forms of cardiac arrhythmias, cardiac alternans feature prominently. This rhythm disturbance at the level of a single cardiac myocyte is characterised by alternating patterns of the membrane potential and the intracellular calcium (Ca^2+^) concentration on successive beats. For instance, at one beat, a long action potential duration (APD) is accompanied by a large intracellular Ca^2+^ transient, while on the next beat, the APD is shortened concomitant with a small amplitude Ca^2+^ transient. As a consequence, contractile efficiency is impaired, which in turn can cause a detrimental reduction in blood flow. In early experimental studies, the intracellular Ca^2+^ concentration was averaged across a cardiac myocyte. The advent of high-resolution microscopy revealed that alternating Ca^2+^ dynamics were already present at individual Ca^2+^ release units (CRU). While one CRU follows a pattern of large-small-large Ca^2+^ transients, neighbouring CRUs exhibit small-large-small Ca^2+^ transients. Crucially, both CRUs experience the same membrane potential. These findings gave rise to the concept of subcellular Ca^2+^ alternans [2–10] and illustrated that nonlinear processes govern cardiac dynamics across multiple scales: the cell wide membrane potential and the Ca^2+^ fluxes restricted to single dyadic clefts.

The existence of subcellular Ca^2+^ alternans reinforces the notion of cardiac myocytes as a network of networks. Each CRU can be conceptualised as a network of interacting components such as L-type Ca^2+^ channels, sodium-calcium exchangers (NCXs) and ryanodine receptors (RyRs). These local networks are then coupled via Ca^2+^ diffusion through both the cytosol and the sarcoplasmic reticulum (SR). This interconnectedness offers multiple explanations for the origin of subcellular Ca^2+^ alternans. On the one hand, we have previously shown that Ca^2+^ alternans can emerge purely through coupling [11]. In other words, an isolated CRU displays a regular period-1 orbit, but upon increasing the coupling strength between CRUs, period-2 orbits characteristic of Ca^2+^ alternans emerge. Crucially, the shape of the Ca^2+^ alternans may depend on the form of coupling. In a recent model of Ca^2+^ cycling, Ca^2+^ alternans emerge for dominant cytosolic coupling via the traditional period-doubling bifurcation, where an eigenvalue of the associated linearised map exits the unit disk through −1 along the real axis [12, 13]. Here, each node exhibits alternating Ca^2+^ dynamics, and neighbouring nodes (or nodes in different parts of a cell) oscillate out-of-phase with each other. For dominant luminal coupling, there is a saddle-node bifurcation, where the leading eigenvalue leaves the unit disk at +1 along the real axis. In this case, each node follows a period-1 orbit, but the amplitudes of neighbouring CRUs varies. On the other hand, changes to the molecular components of a CRU can induce Ca^2+^ alternans, exemplified by weakening sarco-endoplasmic Ca^2+^ ATP (SERCA) pumps or increasing Ca^2+^ flux through L-type Ca^2+^ channels.

To date, investigations on how Ca^2+^ alternans emerge due to modifications at the CRU level have almost exclusively focussed on dominant cytosolic coupling [3, 14–20]. However, the question as to whether Ca^2+^ diffusion in the SR is slow or fast — and hence weak or strong — is still unanswered [21–23]. Here, we focus on strong SR Ca^2+^ diffusion and explore the impact of two modifiers of the local Ca^2+^ dynamics on the genesis of subcellular Ca^2+^ alternans: L-type Ca^2+^ channels and Ca^2+^ buffers.

The L-type Ca^2+^ channel has received significant attention due to its central role in excitation-contraction coupling [24–27]. Its contribution to the formation of Ca^2+^ alternans is ambiguous though [28]. On the one hand, several studies have provided compelling evidence that altering the dynamics of L-type Ca^2+^ channel through e.g. cooperative gating or reducing the current can either promote or inhibit Ca^2+^ alternans [29, 30]. On the other hand, Ca^2+^ alternans have been observed with clamped membrane voltage, thus limiting the degree of control that L-type Ca^2+^ channels can exert on the genesis of Ca^2+^ alternans [31]. Here, we investigate the role of Ca^2+^-dependent inactivation of the L-type Ca^2+^ channel on the dynamics of a CRU, which occurs in addition to voltage-dependent activation and inactivation. [32, 33]. We find that Ca^2+^ -dependent inactivation affects the formation of subcellular Ca^2+^ alternans in a nontrivial manner that depends on the unitary current of the L-type Ca^2+^ channel.

Ca^2+^ buffers are essential for cardiac function, not least because activation of the cytoplasmic buffer troponin C determines how strongly a cardiac myocyte contracts [24, 34]. In addition, the buffers calsequestrin and calmodulin have been shown to vitally shape the dynamics of cardiac myocyte including an impact on the refractoriness of RyRs [35–49]. As has been demonstrated both experimentally and theoretically for numerous cell types and Ca^2+^ releasing channels, including the inositol-1,4,5-trisphosphate receptor, Ca^2+^ buffers can fundamentally alter the dynamics of intracellular Ca^2+^ dynamics ranging from local Ca^2+^ release events such as Ca^2+^ sparks and Ca^2+^ puffs to global Ca^2+^ patterns such as travelling Ca^2+^ waves. Due to the nonlinear dynamics of Ca^2+^ buffers, direct predictions are difficult to make. We show through numerical simulations that Ca^2+^ buffers can both promote and inhibit subcellular Ca^2+^ alternans, which adds another facet to the already rich repertoire of buffered Ca^2+^ dynamics.

## 2. Materials and methods

We consider a two-dimensional network of 15×10 CRUs, where the dynamics of a node with label *μ* is governed by the Shiferaw-Karma model [31]

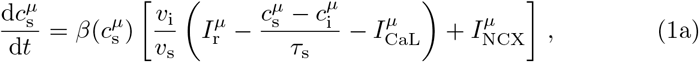

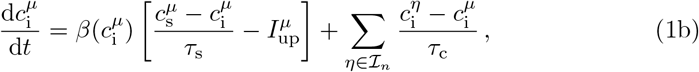

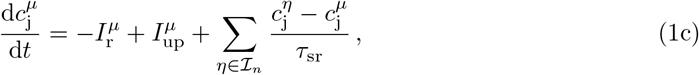

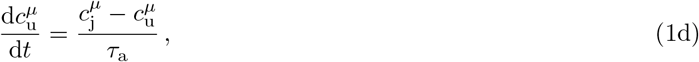

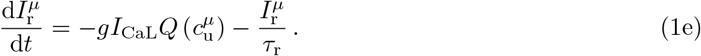

The Ca^2+^ concentrations in the subsarcolemmal space and in the cytosolic bulk are denoted by 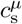 and 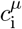, respectively, while the total Ca^2+^ concentration in the SR and the Ca^2+^ concentration in the unrecruited SR are given by 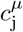 and 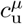, respectively. The Ca^2+^ release current from the unrecruited SR into the subsarcolemmal space is 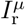, and we refer to the L-type Ca^2+^ current, the NCX current and the SERCA uptake current by 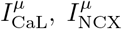 and 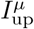, respectively. The model contains four diffusive currents with timescales *τ*_s_, *τ*_c_, *τ*_sr_ and *τ*_a_, describing coupling between the subsarcolemmal space and the cytosolic bulk, through the cytosolic bulk between neighbouring CRUs (indexed by 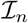), between the total and unrecruited SR, and through the SR between neighbouring CRUs (indexed by 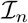), respectively. In some instances, we report the network coupling strengths as inverse of the timescales, i.e. 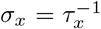, *x* ∈ {c, sr}. The L-type Ca^2+^ current is 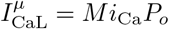, where *M* is the number of L-type Ca^2+^ channels per dyadic cleft, *i*_Ca_ is the single channel current and *P*_*o*_ = *dqf* is the open probability. Here, *d* is the value of the fast voltage-dependent activation gate, *q* corresponds to the Ca^2+^-dependent inactivation gate and *f* to the voltage-dependent inactivation gate. All gates are described by first order kinetics of the form

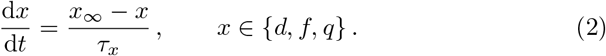

Of particular interest for the present study is

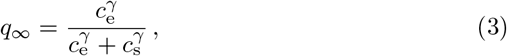

where *c*_e_ sets the *EC*_50_ value, i.e. the value of the subsarcolemmal Ca^2+^ concentration *c*_s_ at which *q*_∞_ equals 0.5, and *γ* controls the sensitivity of Ca^2+^-dependent inactivation. Essentially, the larger *γ* the more step-like the inactivation around a Ca^2+^ concentration of *c*_e_. Buffering is modelled based on the fast-buffer approximation [50, 51] yielding

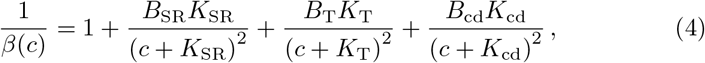

where *B*_SR_ denotes the total buffer concentration in the SR and *K*_SR_ the associated dissociation constant. Constants with the subscript T and cd have the same interpretation, but correspond to troponin C and calmodulin, respectively. For all other details of the model including the functional forms *i*_Ca_ and *I*_NCX_, we refer the reader to [31]. A list of all parameter values used in this study is provided in Table 1. All simulations are performed under clamped voltage conditions, and the we employ no-flux boundary conditions.

**Table 1:**
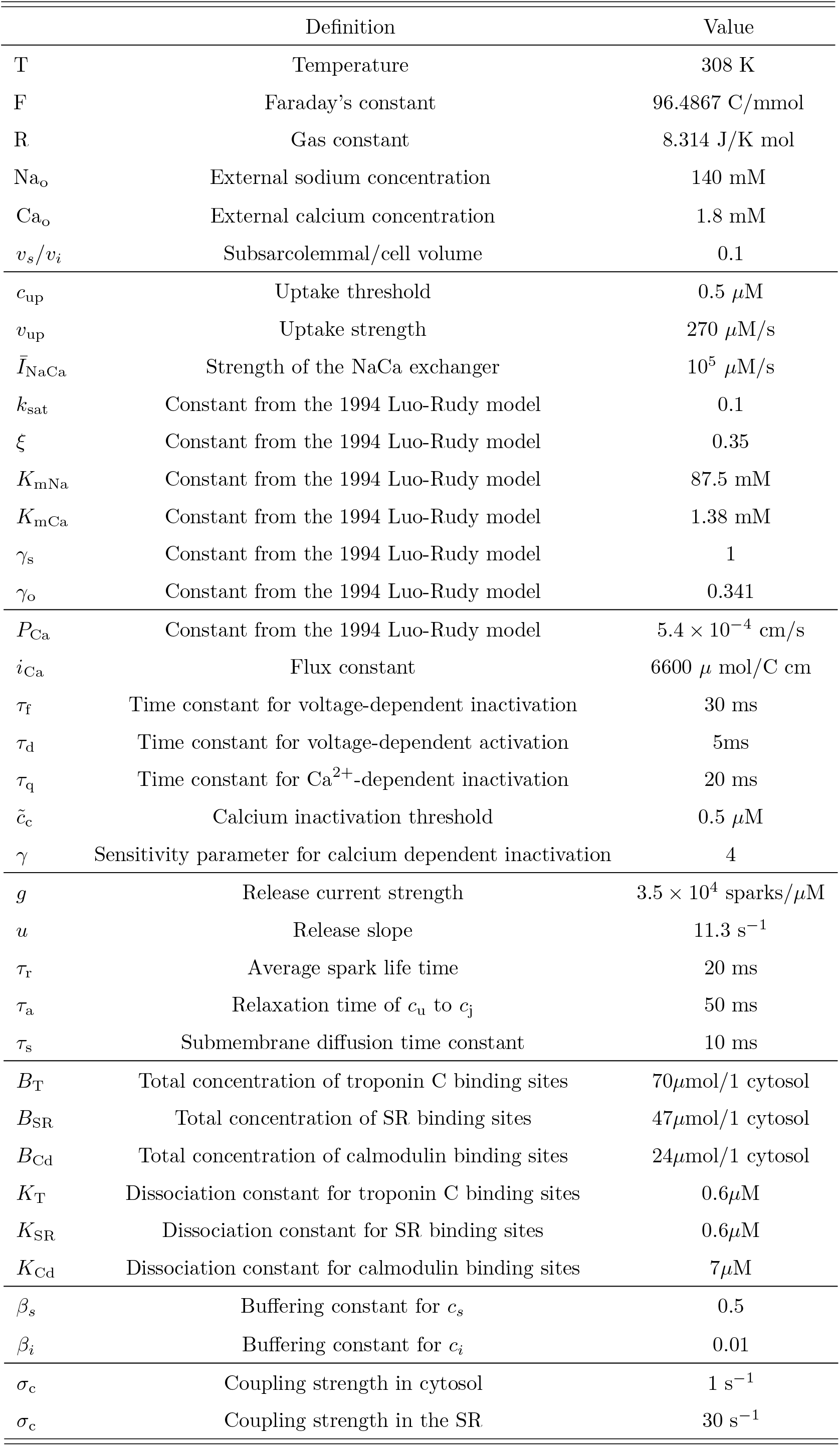
Standard parameter values used in the study.

## 3. Results

To establish a baseline for our findings, we first investigate the dynamics of the CRU network when buffers are clamped over time. In other words, we set 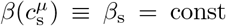 and 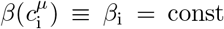 for all *μ*. When cytosolic coupling is dominant, i.e. *τ*_c_ ≪ *τ*_sr_ (*σ*_c_ ≫ *σ*_sr_), synchrony is stable for low pacing frequencies as demonstrated in Fig. 1A. We here show a space-time plot of the *unravelled* CRU network, where we index nodes beginning with 1 in the bottom left corner of the two-dimensional CRU network and move upwards in a row-like manner, i.e. index 16 refers to the most left CRU in the second row from the bottom. When we decrease *T*_*p*_, we observe the emergence of subcellular Ca^2+^ alternans as depicted in Fig. 1B. Each CRU follows a period-2 orbit, where a small amplitude Ca^2+^ transient on one beat is followed by a large Ca^2+^ transient on the next beat. Figures 1C and 1D provide a more detailed view on the emergent spatial pattern, where we plot the peak subsarcolemmal Ca^2+^ concentration on successive beats. The Ca^2+^ alternans are arranged in an *inside-out* pattern along the long axis of the network, where CRUs within one row show almost identical behaviour, but peak amplitudes vary along the vertical direction. When Ca^2+^ transients are large in the centre, they are small towards the top and bottom. On the next beat, this pattern is reversed with large Ca^2+^ transients at the top and bottom.

**Figure 1:**
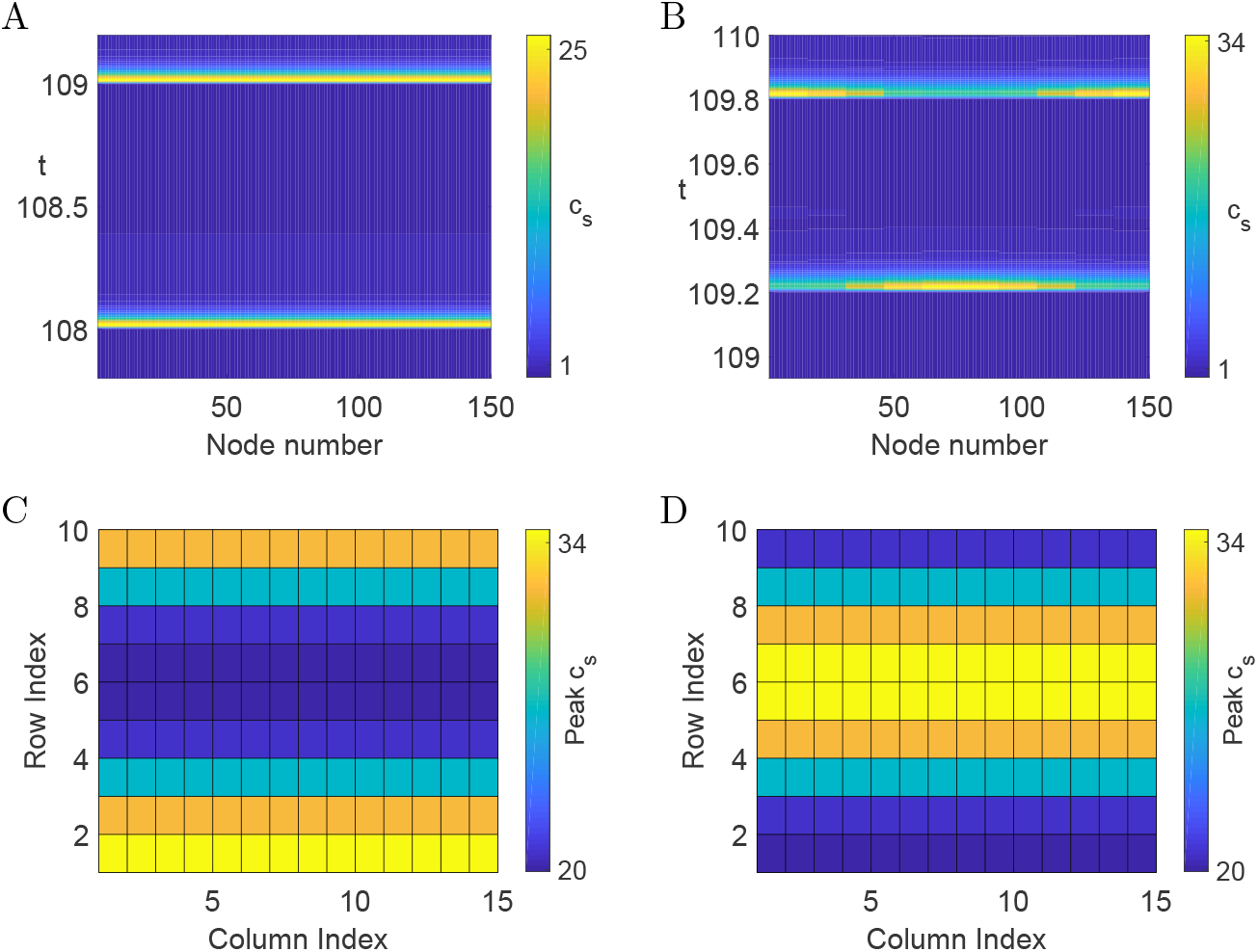
Space-time plot of the subsarcolemmal Ca^2+^ concentration of the unravelled CRU network for(A) *T*_*p*_ = 1s and (B) *T*_*p*_ = 0.6s. (C,D) Peak subsarcolemmal Ca^2+^ concentration on two successive beats across the CRU network. Parameter values as in (B). All other parameter values as in Table 1 and *σ*_c_ = 15s^−1^, *σ*_sr_ = 3s^−1^.

For dominant luminal coupling, where *τ*_c_ ≫ *τ*_sr_ (*σ*_c_ ≪ *σ*_sr_), we again find stable synchrony at low pacing frequencies (see Fig. 2A). Indeed, the space-time plot of the subsarcolemmal Ca^2+^ concentration is identical to the one in Fig. 1A, since when all CRUs exhibit the same behaviour, the coupling terms in Eqs. (1b) and (1c) vanish. The main difference between dominant cytosolic and dominant luminal coupling becomes apparent when we lower *T*_*p*_. For the latter, we find subcellular Ca^2+^ alternans that emerge via a saddle-node bifurcation at the network level, in contrast to a period doubling bifurcation for the former. As Fig. 2B highlights, each CRU follows a period-1 orbit, but this orbit differs amongst the CRUs in the network. Figures 2C and 2D illustrate that here, CRUs on the left form large Ca^2+^ amplitude transients, while the transients are smaller towards the right. In the following we will use Figures 2B to 2D as a reference case and contrast them with the network behaviour when we alter the behaviour of the L-type Ca^2+^ channel and that of Ca^2+^ buffers.

**Figure 2:**
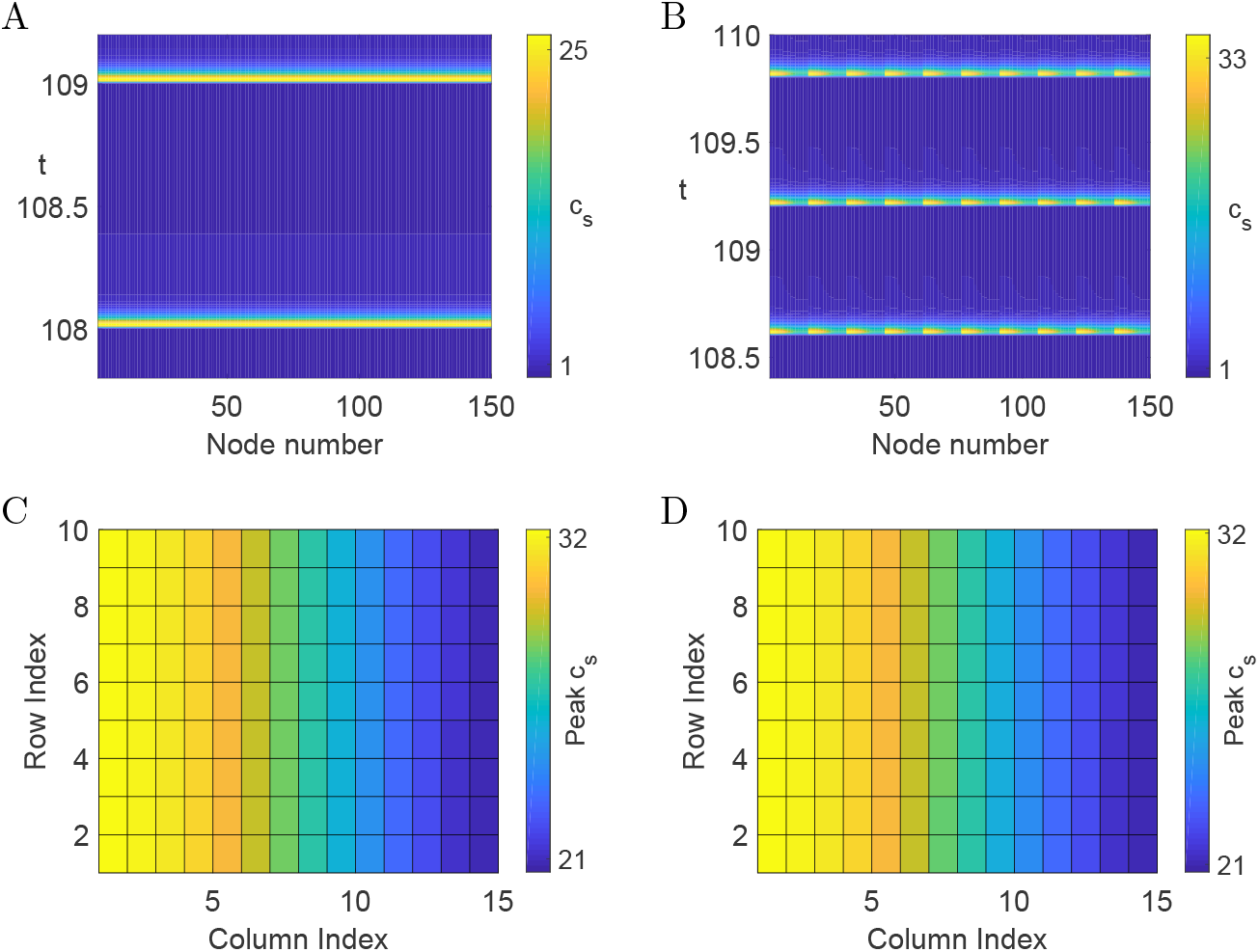
Space-time plot of the subsarcolemmal Ca^2+^ concentration of the unravelled CRU network for(A) *T*_*p*_ = 1s and (B) *T*_*p*_ = 0.6s. (C,D) Peak subsarcolemmal Ca^2+^ concentration on two successive beats across the CRU network. Parameter values as in (B). All other parameter values as in Table 1 and *σ*_c_ = 2s^−1^, *σ*_sr_ = 15s^−1^.

### 3.1. *L-type* Ca^2+^ *channel*

The extent to which Ca^2+^-dependent inactivation sets in as a function of the subsarcolemmal Ca^2+^ concentration 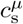 is controlled by the exponent *γ* in Eq. (3). When *γ* is small, the inverse Hill function *q*_∞_ drops slowly from 1 to 0, while a large value of *γ* leads to switch-like behaviour around a concentra-tion value of *c*_e_. As Fig. 3A illustrates, the synchronous network state is stable when *γ* is small. On the other hand, as *γ* is increased, subcellular Ca^2+^ alternans emerge via a saddle-node bifurcation as shown in Figs. 3B-3D. Figure 3B displays a space-time plot of the unravelled CRU network where the variation of the maxima of the Ca^2+^ transients is clearly visible as the colour changes from yellow to blue when we traverse the network. We can also discern changes in the duration of the Ca^2+^ transient as evidenced by the wedge shape of the yellow regions of increased Ca^2+^. Figures 3C and 3D provide more detail on the spatial pattern of the subcellular Ca^2+^ alternans. On each beat, large Ca^2+^ transients occur towards the left side of the myocyte, while Ca^2+^ transients are small towards the right side. Note that there is no variation of the Ca^2+^ peak amplitudes along the row index. These results suggest that a more gradual Ca^2+^-dependent inhibition of the L-type Ca^2+^ channel, i.e. when *γ* is small, protects cardiac myocytes from subcellular Ca^2+^ alternans.

**Figure 3:**
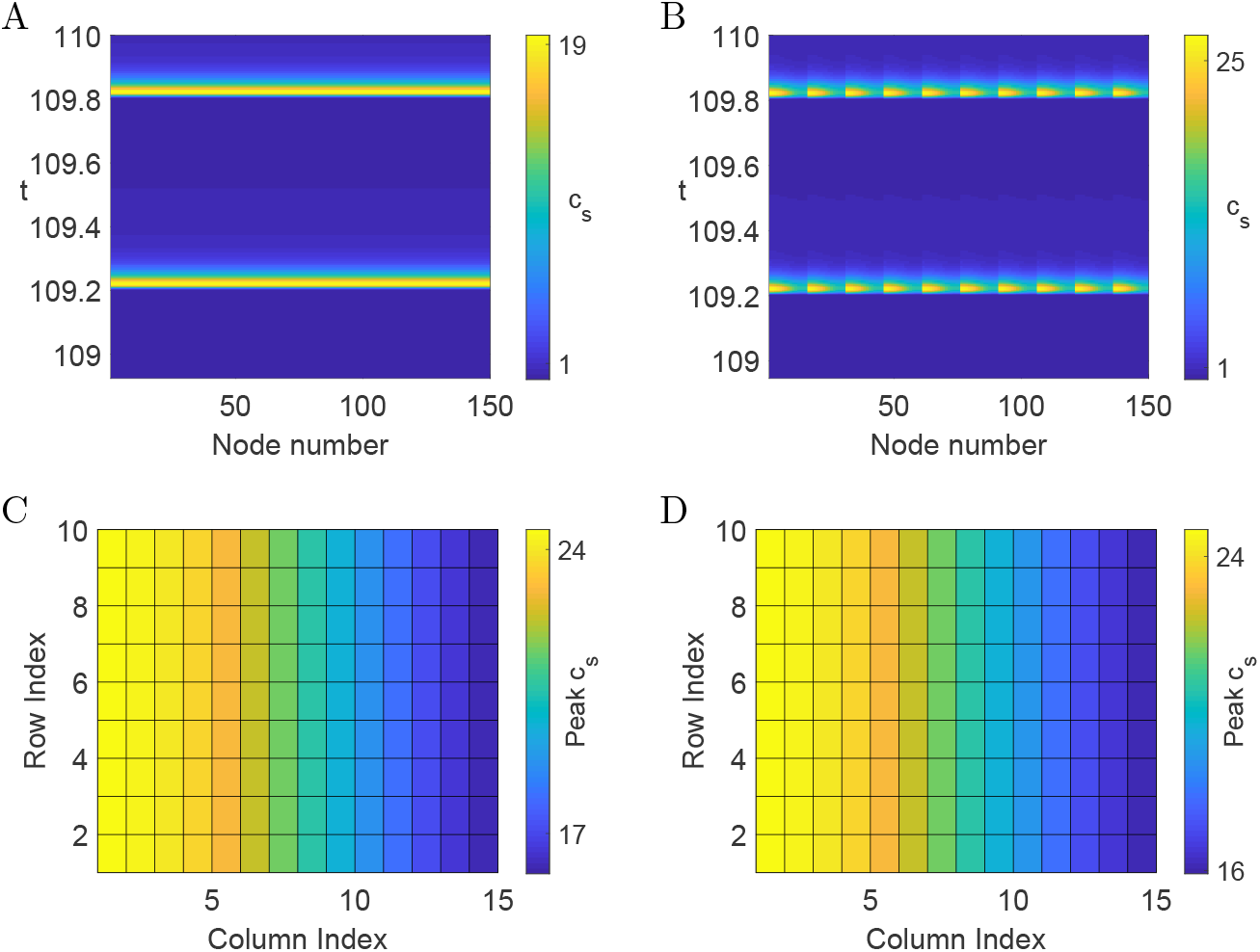
Space-time plot of the subsarcolemmal Ca^2+^ concentration of the unravelled CRU network for *i*_Ca_ = 4400*μ*mol C^−1^cm^−1^ and (A) *γ* = 1, (B) *γ* = 3. (C,D) Peak subsarcolemmal Ca^2+^ concentration on two successive beats across the CRU network. Parameter values as in (B). For all other parameter values, see Table 1.

The unitary current of an L-type Ca^2+^ channel can be modulated through various mechanisms, including *β*-adrenergic stimulation. The space-time plot in Fig. 4A shows that for small values of *i*_Ca_, synchrony is stable. However, upon increasing the single channel current, subcellular Ca^2+^ alternans emerge via a saddle-node bifurcation. We again observe variations of the Ca^2+^ transients in the network similar to those plotted in Fig. 3B. The main difference is the spatial organisation. While the Ca^2+^ transients are most pronounced on the left side in Fig. 3, we find here the largest Ca^2+^ transients towards the right side. This is already discernible in Fig. 4B, where the tip of the yellow wedges points towards the left (in comparison, the yellow wedges point towards the right in Fig. 3B). A clearer view is provided in Figs. 4C and 4D which show the peak subsarcolemmal Ca^2+^ concentration at subsequent beats. The results plotted in Fig. 4 are consistent with experimental findings that upregulation of the L-type Ca^2+^ channel is pro-arrhythmogenic [30].

**Figure 4:**
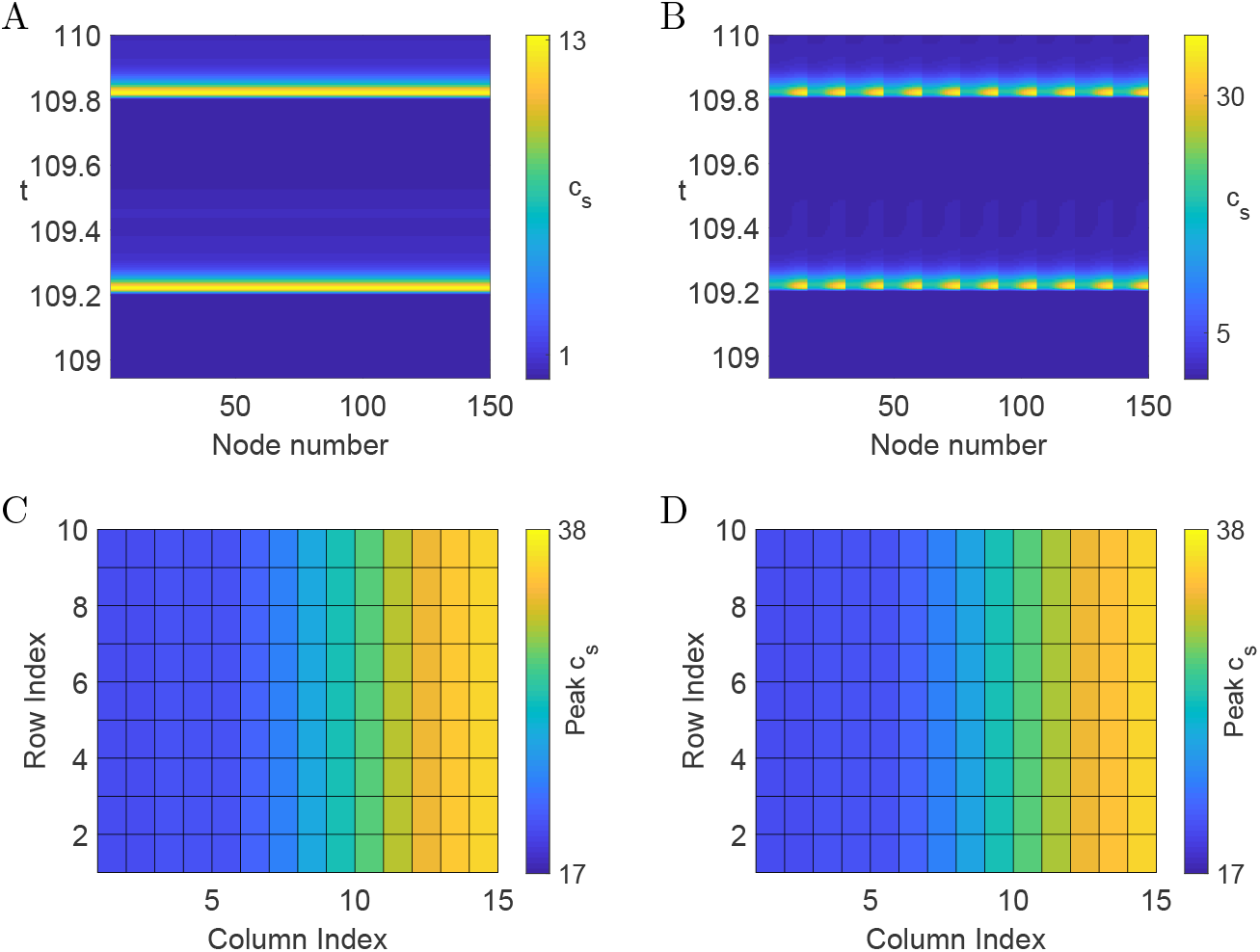
Space-time plot of the subsarcolemmal Ca^2+^ concentration of the unravelled CRU network for *γ* = 3 and (A) *i*_Ca_ = 2200*μ*mol C^−1^cm^−1^, (B) *i*_Ca_ = 6600*μ*mol C^−1^cm^−1^. (C,D) Peak subsarcolemmal Ca^2+^ concentration on two successive beats across the CRU network. Parameter values as in (B). For all other parameter values, see Table 1.

The Ca^2+^ profiles depicted in Figs. 3 and 4 suggest that the effect of the unitary L-type Ca^2+^ current on the generation of subcellular Ca^2+^ alternans depends on the properties of Ca^2+^-dependent inactivation of the channel and vice versa. In Fig. 5 we provide a more comprehensive view on the interplay between these two components. For a given pair of *γ* and *i*_Ca_, we compute the maximal difference in peak subsarcolemmal Ca^2+^ on successive beats for a CRU with index *μ*, i.e

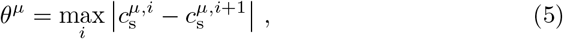

where 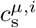 is the maximum of 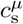 on the *i*th beat. Then, we determine the maximum of all *θ*^*μ*^ across the CRU network, *θ* = max_*μ*_ *θ*^*μ*^. When *i*_Ca_ is small, *θ* vanishes irrespective of the value of *γ*, indicating that synchrony is stable and does not depend on how quickly Ca^2+^-dependent inactivation sets in. For larger values of *i*_Ca_, we observe a sharp transition from synchrony (blue) to alternans (yellow) upon increase of *γ*. Hence, for a sufficiently strong unitary L-type Ca^2+^ current, subcellular Ca^2+^ alternans can be induced if Ca^2+^-dependent inactivation becomes more switch-like. When Ca^2+^-dependent inactivation sets in more gradually, i.e. *γ* is small, synchrony is stable as we increase *i*_Ca_. However, for larger values of *γ*, we observe a sharp transition from synchrony to subcellular Ca^2+^ alternans as the L-type Ca^2+^ channel becomes stronger. There appears to be an L-shape stability boundary in that for a large range of *γ*, subcellular Ca^2+^ alternans appear for approximately the same value of *i*_Ca_, while for a large range of *i*_Ca_, alternans set in for approximately the same small value of *γ*. We also note that the transition from stable synchrony to subcellular Ca^2+^ alternans is quite abrupt, as indicated by the sharp transition from blue to yellow. Taken together, our findings provide strong evidence that the L-type Ca^2+^ channel can initiate subcellular Ca^2+^ alternans, either via its Ca^2+^-dependent inactivation or the strength of its unitary Ca^2+^ current.

**Figure 5:**
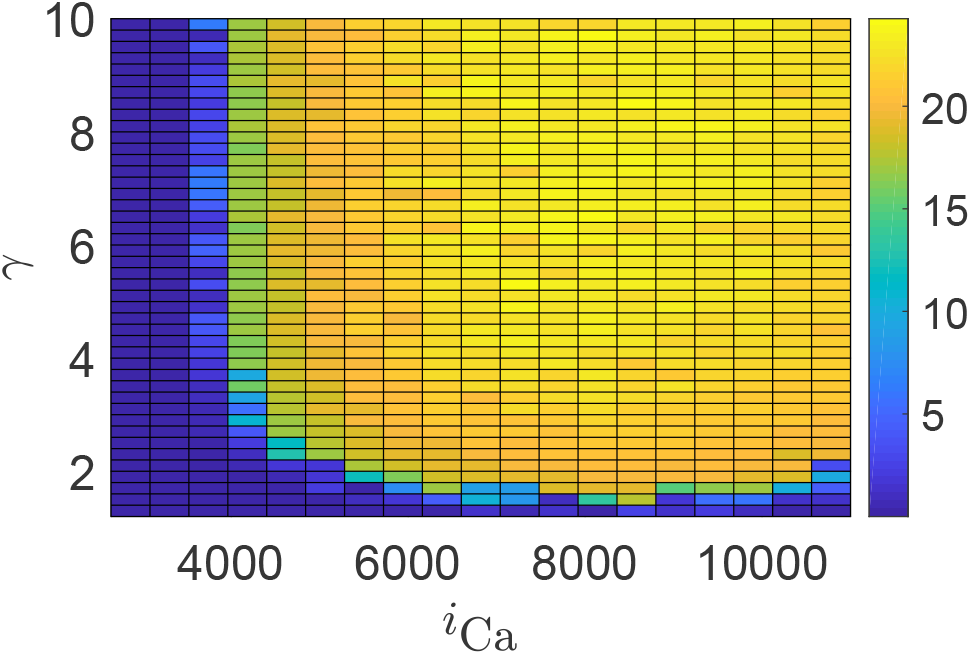
Maximal beat-to-beat variation *θ* of the subsarcolemmal Ca^2+^ concentration as a function of *γ* and *i*_Ca_. All other parameter values as in Table 1.

### 4. Buffers

All results so far were obtained for constant buffer contributions. In other words, we set 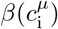 and 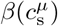 to constants *β*_i_ and *β*_s_, respectively, consistent with earlier work [11]. In this way, we eliminate any time-dependent modulation of the Ca^2+^ dynamics through binding and unbinding to Ca^2+^ buffers. Under more general conditions, however, Eq. (4) entails that 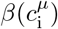 and 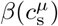 oscillate with the same frequency as 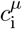 and 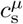, respectively. Figure 6A illustrates that in this case, subcellular Ca^2+^ alternans can be abolished and synchrony is stable. This behaviour needs to be contrasted with that depicted in Fig. 4B, which we would obtain with the parameter values used in Fig. 6A upon replacing the dynamic buffers with the constant buffers used in Fig. 4B. In other words, while the dynamics of the L-type Ca^2+^ channel can induce subcellular Ca^2+^ alternans (as demonstrated in Fig. 4), dynamic Ca^2+^ buffers can rescue this pathological behaviour. This discrepancy between constant and time-dependent buffers prompted us to explore another form of non-responsive buffers. The sensitivity of buffers is usually determined by their dissociation constants, which in the present study are the three constants *K*_SR_, *K*_Cd_ and *K*_T_ in Eq. (4), as well as the corresponding concentration of binding sites *B*_SR_, *B*_Cd_ and *B*_T_. By choosing appropriate values, we can effectively “desensitise” the Ca^2+^ buffers. As Figs. 6B–6D illustrate for the desensitised dynamics, subcellular Ca^2+^ alternans re-emerge consistent with a saddle-node bifurcation. Figure 6B shows a space-time plot of the unravelled CRU network. Each CRU follows a period-1 orbit, which differs both in amplitude and duration of the Ca^2+^ transient across the network, as can be deduced from the variation of the yellow wedges. A more detailed view on the spatial pattern is provided in Figs. 6C and 6D, which depict peak amplitudes of the subsarcolemmal Ca^2+^ concentration on successive beats. Note that although the subcellular Ca^2+^ alternans emerge through a saddle-node bifurcation, the spatial pattern differs from that observed in Figs. 3 and 4. This is consistent with our earlier findings, which have demonstrated a rich pattern space of subcellular Ca^2+^ alternans [12, 13].

**Figure 6:**
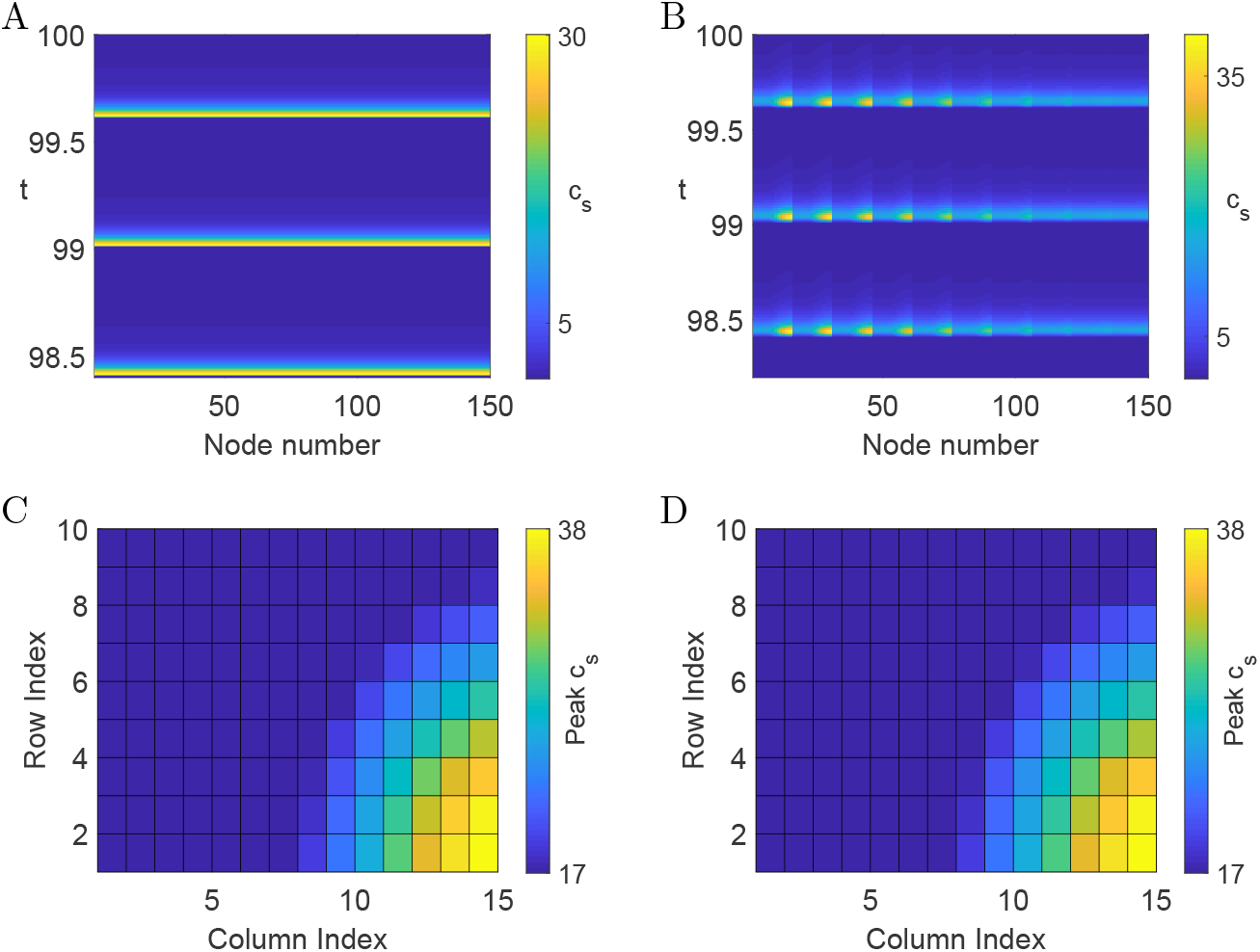
Space-time plot of the subsarcolemmal Ca^2+^ concentration of the unravelled CRU network for (A) fully nonlinear buffers and (B) desensitised buffers (C,D) Peak subsarcolemmal Ca^2+^ concentration on two successive beats across the CRU network. Parameter values as in (B). Other parameter values as in Table 1 and *K*_SR_ = 6.0*μ*M, *K*_T_ = 600.0*μ*M, *K*_Cd_ = 7.0*μ*M, *B*_SR_ = 250.0*μ* mol/1 cytosol, *B*_T_ = 12000.0*μ* mol/1 cytosol, *B*_Cd_ = 1.0*μ* mol/1 cytosol.

At this point, one might be tempted to conclude that constant buffers make the occurrence of subcellular Ca^2+^ alternans more likely. However, as Fig. 7A reveals, this is not the case. Leaving all parameter values unchanged but setting *β*_s_ = *β*_i_ = 1 we find synchrony. Crucially, these simulations correspond to the case without buffers and should be contrasted with the results in Fig. 6A. In both cases, synchrony is stable, but the reasons as to why might differ. The constant values for *β*_s_ and *β*_i_ that we used in Sect. 3.1 were obtained for a piecewise linear (PWL) caricature of the model given by Eq. (1), see [11] for details. To obtain estimates that are more consistent with the full nonlinear model, we determine the mean values of 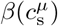 and 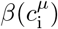 when synchrony is stable and assign them to *β*_s_ and *β*_i_, respectively. For these values, we find subcellular Ca^2+^ alternans that are consistent with a saddle-node bifurcation as shown in Figs. 7B. Again, individual CRUs display a period-1 orbit, which differs throughout the network. The spatial pattern of the subcellular Ca^2+^ alternans is reminiscent of the one depicted in Figs. 3B – 3D, where Ca^2+^ transients are more pronounced on the left side of the CRU network compared to the right side.

**Figure 7:**
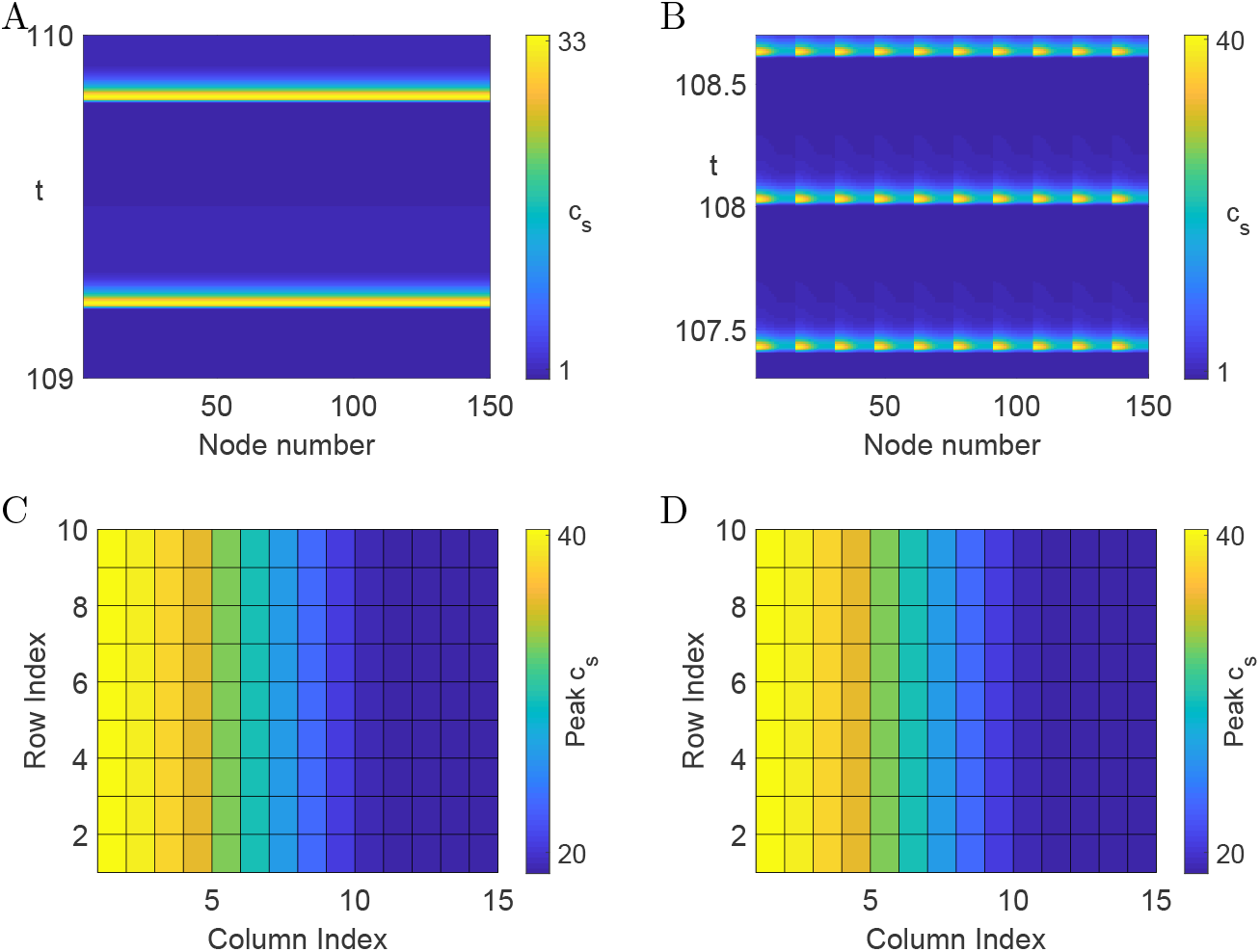
Space-time plot of the subsarcolemmal Ca^2+^ concentration of the unravelled CRU network for *T*_*p*_ = 0.6s and (A) *β*_s_ = *β*_i_ = 1, (B) *β*_s_ = 0.08827, *β*_i_ = 0.01738. (C,D) Peak subsarcolemmal Ca^2+^ concentration on two successive beats across the CRU network. Parameter values as in (B). For all other parameter values, see Table 1.

Since constant Ca^2+^ buffers can both promote as well as abolish subcellular Ca^2+^ alternans, we next explore the impact of the buffer time course on the formation of subcellular Ca^2+^ alternans. To do this in a controlled fashion, we extract the time course of both 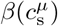 and 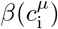 from the full nonlinear model and then clamp the buffer time courses at each node to these profiles. In other words, each node experiences nonlinear buffer dynamics, but the buffers do not alternate from node to node. As Fig. 8A reveals, we obtain subcellular Ca^2+^ alternans that differ from those reported so far in this study. Here, every node in the network follows the same period-2 orbit characteristic of subcellular Ca^2+^ alternans that emerge via a period-doubling bifurcation. Figures 8B and 8C illustrate the uniform behaviour across the network that alternates between successive beats. This spatial pattern is known as spatially concordant Ca^2+^ alternans. When we change the coupling strengths, but keep all other parameter values unaltered, we observe spatially discordant Ca^2+^ alternans as plotted in Figs. 8D–8F. Every node follows a period-2 orbit, but different parts of the network oscillate out-of-phase with each other.

**Figure 8:**
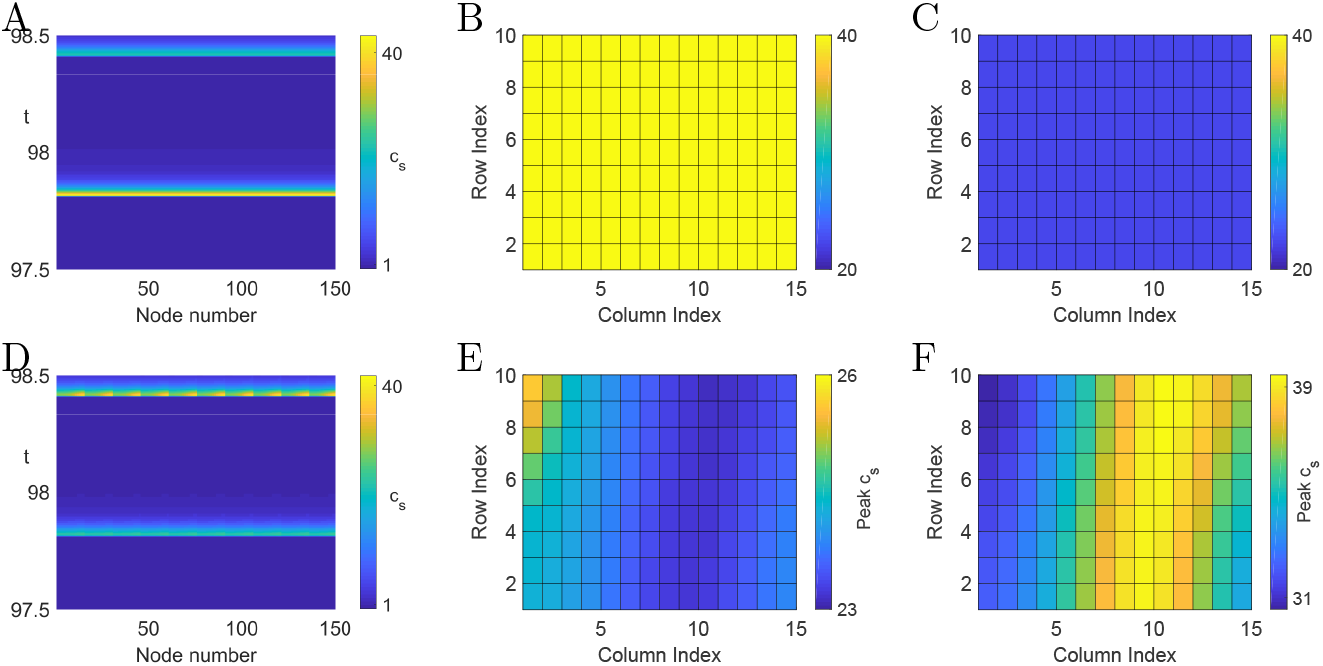
Network dynamics for *T*_*p*_ = 0.6s and clamped nonlinear buffers for (A–C) *σ*_sr_ = 30s^−1^, *σ*_c_ = 1s^−1^, (D–F) *σ*_sr_ = 3s^−1^, *σ*_c_ = 0s^−1^. Space-time plots of the subsarcolemmal Ca^2+^ concentration of the unravelled CRU are shown in (A) and (D). Peak subsarcolemmal Ca^2+^ concentration on two successive beats across the CRU network are plotted in (B,C) and (E,F). For all other parameter values, see Table 1.

Our results so far strongly suggest that the time course and amplitude of Ca^2+^ buffers significantly impacts on the genesis of subcellular Ca^2+^ alternans. Figure 9 shows results from an *in silico* experiment in which we tune the Ca^2+^ buffer dynamics from constant (*ε* = 0) to fully nonlinear (*ε* = 1). As a measure for the strength of subcellular Ca^2+^ alternans, we report the maximal beat-to-beat variation *θ* as defined after Eq. (5). As *ε* increases, we find a monotonic decrease in *θ*, highlighting that nonlinear buffers have the potential to abolish subcellular Ca^2+^ alternans.

**Figure 9:**
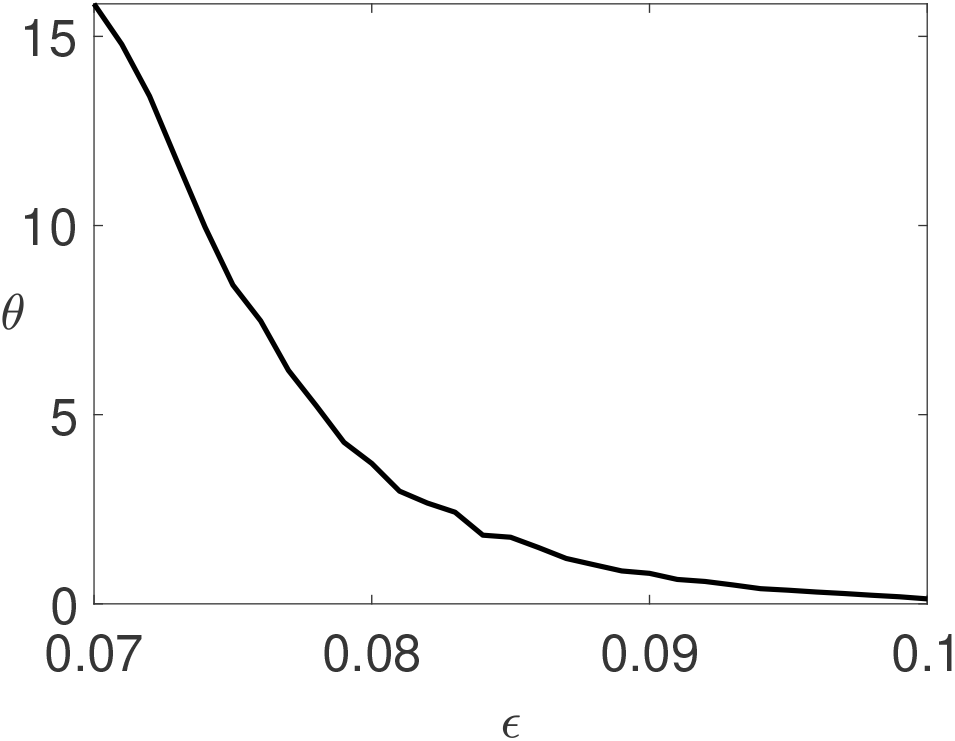
Maximal beat-to-beat variation *θ* of the subsarcolemmal Ca^2+^ concentration as a function of the variability of the Ca^2+^ buffer *ε*. See text for details. Parameter values as in Table 1.

## 5. Discussion

Subcellular Ca^2+^ alternans have been firmly linked to the genesis of cardiac arrhythmias. Despite this crucial connection, we still lack a complete picture of how the dynamics of the intracellular Ca^2+^ concentration transitions from its healthy period-1 orbit to its various pathological forms.

Our focus has been on understanding subcellular Ca^2+^ alternans in tubulated myocytes, such as ventricular myocytes. The presence of t-tubules in these cells gives rise to well-defined CRUs, which form a network where nearest neighbours are coupled via Ca^2+^ diffusion, both through the cytosol and the SR. The discussion of whether Ca^2+^ diffusion in the SR is fast or slow has been ongoing for more than a decade [21–23], without a resolution in sight. We illustrate in Figs. 1 and 2 that whether Ca^2+^ diffuses more dominantly in the lumen or in the cytosol has major consequences for the spatial patterns of subcellular Ca^2+^ alternans. In the latter, subcellular Ca^2+^ alternans emerge via the classical period-doubling bifurcation, where CRUs exhibit a period-2 orbit and CRUs in different parts of the cell oscillate out-of-phase with each other. This behaviour has been well studied and documented [2–9]. On the other hand, when Ca^2+^ diffusion in the SR dominates, we observe a completely different spatial pattern originating from a saddle-node bifurcation. Here, CRUs show a period-1 orbit, which is different from the synchronous network state and where CRUs in different regions of the cell exhibit Ca^2+^ transients of varying amplitude. It is worth noting that the discussion of whether intraluminal Ca^2+^ diffusion is faster than cytosolic Ca^2+^ diffusion — a process known as intraluminal tunnelling — has already received attention, although in a different context [52]. Given the largely unexplored nature of the saddle-node bifurcation in the generation of subcellular Ca^2+^ alternans, we have concentrated on dominant luminal coupling in the present study and have investigated two main contributors that shape the dynamics of cardiac Ca^2+^: the L-type Ca^2+^ channel and Ca^2+^ buffers.

The L-type Ca^2+^ channel constitutes a major Ca^2+^ conduit that regulates Ca^2+^ influx from the extracellular space into the myoplasm and is thus crucial for high-fidelity excitation-contraction coupling. It is therefore not surprising that pathologies of the L-type Ca^2+^ channel can lead to abnormal Ca^2+^ dynamics. When we increase the single channel Ca^2+^ current *i*_Ca_, subcellular Ca^2+^ alternans are more likely to occur as evidenced by the transition from blue to yellow in Fig. 5. However, this behaviour depends on the strength of Ca^2+^ -dependent inactivation of the L-type Ca^2+^ channel. As is often the case, the inactivation gate is modelled via a first-order kinetic scheme with a time constant *τ_q_* and a state-dependent steady state *q*_∞_. As Eq. (3) shows, *q*_∞_ follows an inverse Hill function with exponent *γ*. Hence, for small values of *γ*, *q*_∞_ changes gradually as a function of the subsarolemmal Ca^2+^ concentration *c*_s_. On the other hand, large values of *γ* lead to a switch-like Hill function. When *i*_Ca_ is small, the increase in subsarolemmal Ca^2+^ is small as well, which in turn almost completely eliminates Ca^2+^ dependent inactivation (as *q* never falls sufficiently towards zero). Therefore, we do not observe any effect of *γ* on the generation of subcellular Ca^2+^ alternans in this regime, indicated by the blue band towards the left of Fig. 5. On the other hand, as we increase *i*_Ca_, the larger subsar-colemmal Ca^2+^ concentrations allow for a larger exploration of the right tail of *q*_∞_, and hence values closer to zero. When *γ* is large making *q*_∞_ more steplike, bigger values of 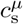 entail longer periods where *q* tends to zero. An increase of *i*_Ca_ does not change that, meaning that the nature of the subcellular Ca^2+^ alternans is not affected by increasing *i*_Ca_ for larger values of *γ*. This explains the almost uniform yellow colouring in Fig. 5 for fixed large *γ* and varying *i*_Ca_. An interesting feature of Fig. 5 is the sharp transition from regular behaviour to subcellular Ca^2+^ alternans as is manifest from the abrupt colour change from blue to yellow. It remains to be seen whether this behaviour can be understood more formally in terms of a phase transition.

All results for the L-type Ca^2+^ channel were obtained with constant buffer contributions. However, since the concentration of Ca^2+^ bound buffers directly depends on the intracellular Ca^2+^ concentration, the buffer function *β* in Eq. (4) should evolve over time. Using the full nonlinear buffers, we find that subcellular Ca^2+^ alternans are extinguished (Fig. 6A). In other words, solely changing the buffer dynamics completely alters the dynamics of the cardiac cell. These findings are in line with a large body of literature demonstrating that Ca^2+^ buffers can substantially modify intracellular Ca^2+^ dynamics. From a physiological perspective, our results indicate that Ca^2+^ buffers can perform a stabilising role that can compensate for dysfunctions of other components of the Ca^2+^ signalling toolkit, such as the L-type Ca^2+^ channel. Because Ca^2+^ buffers are slaved to the Ca^2+^ dynamics, the buffer dynamics exhibit alternans as soon as the intracellular Ca^2+^ concentration alternates. For the results in Fig. 8, we broke this connection and clamped the Ca^2+^ buffer dynamics in such a way that each node exhibits the same nonlinear orbit. In other words, Ca^2+^ buffers alternate at each node, but there is no spatial variation of the buffer dynamics. In this regime, the patterns of the subcellular Ca^2+^ alternans vary drastically from the ones we observed so far. We found spatially concordant alternans, which can transition into spatially discordant alternans upon altering the coupling strength of cytosolic and SR diffusion. While we employed buffers to induce this pattern change, it is conceivable that such dynamics could originate from other dynamical variables of cardiac Ca^2+^ cycling. In this case, our results point to more subtle dependencies in that the nonlinear dynamics of cardiac Ca^2+^ cycling can be easily disturbed into new dynamic regimes, potentially inducing a plethora of cardiac arrhythmias. It is therefore astonishing that cardiac Ca^2+^ dynamics more often than not behaves completely regularly; a fact that certainly deserves more attention.

We first reported the emergence of subcellular Ca^2+^ alternans via a saddle-node bifurcation in a PWL caricature of an established Ca^2+^ cycling model [12]. One might wonder if this novel form of subcellular Ca^2+^ alternans is a consequence of the approximations used in the derivation of the PWL model. The results presented here show that this is not the case. The fully nonlinear model exhibits the same instabilities. This provides further evidence that PWL models are valuable in exploring the behaviour of complex nonlinear systems and thus adds to earlier success stories such as the McKean model, which represents a PWL version of the Fitzugh-Nagumo model for the propagation of neural action potentials [53–55]. The advantage of PWL models is that the majority of the analysis can be performed semi-analytically, which greatly facilitates the exploration of the associated parameter space. In turn, this allows for a more comprehensive classification of the possible dynamics. In contrast, fully nonlinear systems can often only be dissected via direct numerical simulations, which is often only done for a small subset of parameter values. In this respect, PWL models can provide guidance for the analysis of the nonlinear systems and where to explore in parameter space for interesting behaviour.

The last point becomes especially pertinent for the exploration of the different spatial patterns that emerge via a saddle-node bifurcation. As Figs. 1, 2, 6 and 8 illustrate, the Ca^2+^ profiles across the network exhibit significant variability. In a PWL model, these patterns can be classified and understood from a linear stability analysis, which can be performed in closed form [12, 13]. On the other hand, the nonlinear model requires direct simulations, which are computationally more expensive and limited in scope as to what parameter values to sample.

As stated above, our focus here is on tubulated myocytes. However, Ca^2+^ alternans have also been observed in non-tubulated cells such as atrial myocytes and failing ventricular myocytes [56–62]. In these cells, L-type Ca^2+^ channels are only located at the cell periphery, where they trigger Ca^2+^ release from the SR through the RyR. A Ca^2+^ wave then propagates centripetally from the periphery via diffusion and Ca^2+^ induced Ca^2+^ release [63, 64]. Conceptually, it therefore makes sense to distinguish junctional CRUs (that contain L-type Ca^2+^ channels) and non-junctional CRUs (that lack L-type Ca^2+^ channels). Due to the stronger reliance on Ca^2+^ diffusion, it will be interesting to explore how differences in the diffusive coupling between CRUs and the fact there are two classes of CRUs shape subcellular Ca^2+^ alternans and whether the bifurcation structure observed for tubulated myocytes carries over to non-tubulated ones. Answering this question will not only unravel further similarities or differences between tubulated and non-tubulated myocytes, it will also advance our understanding of atrial fibrillation, which is projected to become epidemic with an ageing population [65].

## Acknowledgement

This work was supported by the Engineering and Physical Sciences Research Council [grant number EP/P007031/1].

## Appendix

We here provide the parameter values used in the study unless otherwise stated.

